# The multi-TLR inhibitor soluble TLR2 suppresses Chronic Kidney Disease-induced exacerbation of atherosclerosis by inhibiting DAMPs activity

**DOI:** 10.64898/2026.11.24.769056

**Authors:** Esra Cetin, Alex Gibbs, Asmita Chaudhari, Sumukh Deshpande, Robert Andrews, Soma Meran, Dipak P. Ramji, Donald Fraser, Mario O. Labéta, Anne-Catherine Raby

## Abstract

**Objective:** Chronic kidney disease (CKD) markedly increases cardiovascular (CV) risk, with exacerbated atherosclerosis as a major contributing factor. CKD-associated chronic inflammation is believed to drive this association, however, the specific mechanisms remain to be defined and, thus, therapeutic options remain limited. This study aims to establish the role of the damage-associated molecular pattern (DAMP):Toll-like receptor (TLR) signaling axis in driving atherosclerosis in CKD and to evaluate DAMP:TLR pathway-targeting therapies to alleviate CKD’s worsening effect.

**Approach and Results:** Chronic nephropathy in mice replicated key cardiovascular features observed in CKD patients, including: i) elevated plasma inflammatory cytokines, ii) increased circulating inflammatory monocyte proportions; iii) bigger atherosclerotic lesions that were iv) more inflammatory with a more vulnerable phenotype. In support of a role for the DAMP: TLR pathway in driving these effects, nephropathic mice displayed increased circulating levels of several DAMPs and CKD patients’ proteomic data highlighted multiple DAMPs also elevated in plasma from patients who later experienced cardiovascular events. By conducting the first single-cell transcriptomic analysis of murine atherosclerotic aortas in chronic nephropathy, enrichment of TLR-related pathways across vascular and immune cell populations was revealed.

Although Calprotectin was the most elevated DAMP in CKD patients who later underwent a CV event and was also elevated in nephropathic mice, its inhibition only modestly improved atherosclerosis. Conversely, treatment with soluble TLR2, a multiple TLR inhibitor, robustly counteracted the atherosclerosis-exacerbating effects of chronic nephropathy, confirming a role for the DAMP:TLR pathway in driving the observed pathology and suggesting that inhibition of multiple DAMPs may be required to ameliorate the observed pathology.

**Conclusion:** Thus, this study provides the foundation for future clinical studies aimed at evaluating soluble TLR2 as a therapeutic agent to improve cardiovascular outcomes in this challenging patient population, demonstrating efficacy in established chronic nephropathy while critically preserving host antibacterial defence.

## INTRODUCTION

Increased risk of cardiovascular (CV) mortality in CKD can equate to up to 20 years of life lost^1,2^. Notably, CKD patients typically display an increased burden of atherosclerosis^2–4^, an inflammatory condition of the arterial wall that underlies most CV events^5,6^. Traditional risk factors do not account for the higher atherosclerotic burden in CKD, and clinical interventions successful in the general population are less effective^2,7^in CKD. Chronic inflammation induced by CKD has been suggested to play a role in promoting CV disease^1,2^, but the detailed mechanisms have not been defined.

Critical to the development of chronic inflammatory pathologies, including atherosclerosis, are the Toll-like receptors (TLRs)^8,9^. TLRs are early mediators of inflammation following infection or tissue damage, the latter via recognition of Damage-Associated Molecular Patterns (DAMPs), endogenous ligands that are released following cell death and tissue injury^10^. We and others have identified specific DAMPs that are elevated in CKD patients and promote chronic inflammation^11,12^. We hypothesise that elevated DAMPs contribute to worsening atherosclerotic disease in the CKD population.

To test this, chronic nephropathy was induced with the nephrotoxin Aristolochic-Acid (AA)^11,13,14^ in atherosclerosis-prone Low-Density Lipoprotein (LDL) Receptor knock-out (*Ldlr*^−/−^) mice. Nephropathic mice displayed key CV/atherosclerotic features observed in CKD patients, including: i) elevated plasma inflammatory cytokines, ii) increased circulating monocyte proportions^15^, iii) bigger atherosclerotic lesions iv) with a more vulnerable phenotype^3^. Supporting a role for DAMPs in mediating these effects, multiple DAMPs were elevated in AAN mice. Human proteomics data also showed several DAMPs at higher systemic concentrations in CKD patients who subsequently experienced a CV event. Consistent with this, single-cell transcriptomic analysis of atherosclerotic aortas - the first conducted in chronic nephropathy - revealed enrichment of pathways in line with TLR activation or downstream of it, in most cell clusters identified.

While Calprotectin was the most elevated DAMP in CKD patients who later underwent a CV event and was elevated in plasma from nephropathic mice, its inhibition only modestly improved atherosclerosis in these animals. Conversely, treatment with the multi-TLR inhibitor soluble TLR2 (sTLR2)^16^ abrogated most of chronic nephropathy’s exacerbating effects on atherosclerotic disease and inhibited nephropathy-elevated transcriptomic pathways across most aortic cell clusters.

These findings reveal the role of the DAMP:TLR pathway in worsening atherosclerosis in CKD and suggest the contribution of multiple DAMPs to the pathology. Critically, this study highlights the efficacy of soluble TLR2 as a therapeutic agent to improve cardiovascular outcomes in this challenging patient population, with the notable advantages that treatment was effective when initiated after chronic nephropathy was established and does not compromise bacterial clearance.

## METHODS

Details in Sup. Methods

### Animal experiments

All procedures were approved by Cardiff University Animal Welfare Ethical Review Body, carried out under a UK Home Office Licence (PP1959620) and conformed to the guidelines from Directive 2010/63/EU of the European Parliament. 8-week-old *Ldlr* ^−/−^ mice (homozygous for the *Ldlrtm1Her* mutation and backcrossed to the C57BL/6J strain for 10 generations, n=11-25/group) received i.p. injections of Aristolochic Acid (AA, 3mg/kg), every 2 days for a total of 6 times, to induce AA-Nephropathy (AAN group), or PBS injections (Control group). After the last injection (Day 12), AAN and Control mice were put on a high-fat diet (HFD, 21% w/w pork lard, 0.15% w/w cholesterol) for 15 weeks (Sup. Fig.1A). For experiments evaluating therapeutic strategies, administration of Paquinimod in drinking water (5mg/kg/day^17^), or weekly i.p. injections of sTLR2 (12.5 µg/kg^11^) were initiated from Day 29. Age-matched *Ldlr*^−/−^ mice not injected and receiving a chow diet served as a “no-atherosclerosis” reference group (Naïve, n=6-8). At the end of the experiment, all animals were euthanised by exposure to a rising gradient of carbon dioxide (0.04 to 70%) followed by total exsanguination (cardiac puncture).

### Kidney fibrosis

Kidney fibrosis was quantified by Masson’s trichrome staining^11^.

### Flow cytometry

Flow cytometric analysis of whole blood was as previously described^11^. Neutrophils were defined as Ly6G^+^ and CD11b^+^ and monocytes as Ly6G^-^ and CD11b^+^.

### Atherosclerotic Lesion Analysis

Frozen sections of the aortic root were obtained and sections displaying the 3 valves or sequential sections (100 µm apart) from the 3 open valves were analysed for atherosclerosis. Plaque size, lipid content and vulnerability phenotype were then assessed by Oil Red O (ORO) staining counterstained with hematoxylin, Macrophage and Smooth Muscle Cell (SMC) composition and distribution were determined by immunofluorescence (CD68 and Smooth Muscle Actin, SMA, respectively).

Analyses were performed in a blinded manner. Plaque size is shown as percentage (%) of occlusion, the ratio between plaque area and vessel lumen area. Plaque lipid content is the ratio between ORO positive area and total plaque area. Necrotic cores were identified as areas devoid of hematoxylin stain within an area of atherosclerotic plaque and fibrotic cap thickness as the shortest distance between the top of the core and the lumen. Animals displaying no plaques with a visible necrotic core (0% necrotic core area) were excluded from the fibrotic cap thickness analysis.

### Plasma markers

Plasma concentration of cystatin, DAMPs, cytokines, OxLDL and endotoxin (LPS) were measured by ELISA. Urea was quantified using a Jung’s method-based assay (Urea Assay Kit II, Abcam). Total cholesterol, LDL/VLDL, and HDL levels were determined using the Cholesterol Assay Kit - HDL and LDL/VLDL (Abcam).

### Single-cell preparation

Thoracic aortas underwent enzymatic digestion for single-cell isolation. Viability was 78-90%. Approximately 500, 000 to 1-million cells/sample were fixed using the Evercode™ Cell Fixation Kit v3 (Parse Biosciences) and kept at -80°C until library preparation.

### Library preparation

Sequencing libraries were prepared from thawed single-cell suspensions using Parse’s Evercode WT Kits v3. In brief, 6250 cells per animal (n=6 animals per experimental group, 3 females, 3 males) went through “split-pool” barcoding steps before being split into sub-libraries. Ten sub-libraries were sequenced on NovaSeq X 25B PE150 flowcells to yield on average 52,000 reads per cell.

### Sequencing data processing and clustering

Raw sequencing reads were processed for: barcode correction, read alignment, read deduplication, transcript quantification, and subsequent generation of cell-by-gene count matrices (Parse Biosciences Trailmaker). Count matrices were then imported into R and Seurat objects were made for each sample using *Seurat* functions^18^ and filtered to contain < 10% mitochondrial genes, and > 200 and < 10,000 detected genes. Filtered samples were then merged into one object and *SCTransform* (with mitochondrial gene regression^19^) was used to normalise, scale, and obtain highly variable genes. Linear dimensional reduction was then performed via principal component analysis with default parameters (50 principal components) before the removal of doublets via the *DoubletFinder* package^20^. *SCTransform* and linear dimensional reduction was again performed on the singlet-containing dataset, followed by batch correction via the *Harmony* package^21^. A nearest-neighbours graph was used to perform low-resolution clustering on the dataset (resolution=0.1). Non-linear dimensional reduction (Uniform Manifold Approximation and Projection, UMAP) was performed, and gene expressions were visualised via *FeaturePlots* for cell annotation.

### Differentially Expressed Genes and Gene Ontology Pathway Analysis

Differentially Expressed Genes (DEGs, Adjusted *p* <0.05 and |fold change|>1.5)) for each cluster were generated via Seurat’s *FindMarkers* function.

Gene ontology enrichment analysis for Kyoto Encyclopedia of Genes and Genomes (KEGG) pathways was performed (ShinyGO) with upregulated and downregulated DEGs separately^22^. Statistical enrichment was determined using a hypergeometric test, with the resulting raw p-values adjusted for multiple testing via the Benjamini-Hochberg false discovery rate (FDR). Enriched KEGG pathways with a False-Discovery Rate (FDR) < 0.05 and a minimum pathway size of 10 were selected. Pathways containing irrelevant terms and pathways enriched because of one gene only were removed from the analysis.

### CKD patients’ proteomics data

Data was repurposed from ^17^ and all methods can be found in the original article. Briefly, full proteomics analysis was performed in serum from 66 patients with CKD stage 3 and 4 (15 < eGFR < 60 ml/min per 1.73 m^2^), including 32 cases who did present with a CV event within the follow-up period (mean, 3.3 ± 1.2 years) and 34 matched controls (age, sex, eGFR, diabetes, and cholesterol) who did not. *p* values are shown before and after adjustment for multiple testing using the Benjamini-Hochberg procedure.

### Statistical analysis

All statistical analyses were performed using GraphPad Prism (10.4.1). Normal (Gaussian) distribution was assessed by Shapiro-Wilk test. All comparisons involved at least one group where normal distribution was not achieved. For experiments comparing 2 groups, *p* values were calculated using a Mann-Whitney U test for independent samples. When comparing 3 or more groups, a Kruskal-Wallis test was used. To determine the effect of sex and its interaction with nephropathy, a two-way ANOVA was performed. Differences with *p* values <0.05 were considered statistically significant. Trends (0.05 <*p* <0.1) are also indicated (numerical *p*).

## RESULTS

### AA nephropathy (AAN) leads to higher and more phenotypically vulnerable atherosclerotic plaque burden

Repeated administration of AA (Sup. Fig.1A) leads to kidney inflammation^11,13,14^, progressing to marked kidney fibrosis (Sup. Fig.1B), kidney atrophy (Sup. Fig.1C), and to a decline in kidney function (increased plasma Cystatin C and Urea, Sup. Fig.1D). We verified that both male and female mice displayed kidney fibrosis and loss of function. Interestingly, urea levels were not elevated when looking at both sexes together, likely due to the effect of sex on urea levels (Fig. 1D, table), but was increased upon nephropathy when looking a sexes separately.

**Figure 1.**
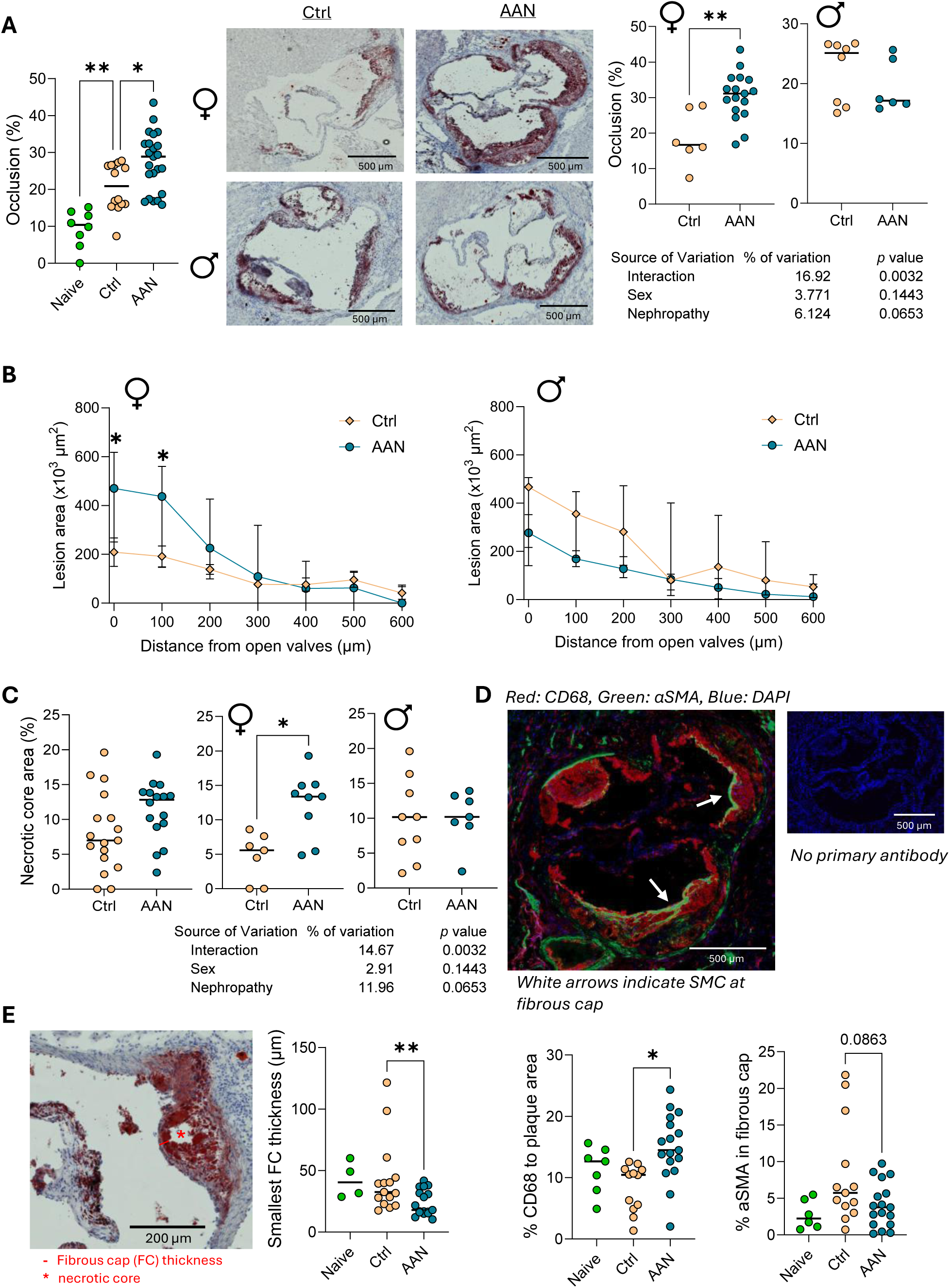
Increased atherosclerotic plaque burden and instability in *Ldlr ^-/-^* mice with AA-induced nephropathy. *Ldlr ^-/-^* male and female mice were injected intraperitoneally 6 times at 2-day intervals with AA (3 mg/kg, AAN) or PBS (Ctrl), starting on Day 0. On Day 12, normal chow diet was replaced with a high-fat diet (HFD) for 15 weeks. Age-matched *Ldlr ^-/-^* mice having not received injections or HFD served as a normal reference group (Naïve). Frozen sections of the aortic root were obtained and sections displaying the 3 valves (A, C-E) or sequential sections (B) were analysed for atherosclerosis. **A-B**. Oil Red O staining was used to determine the percentage of occlusion (plaque to lumen area, representative images shown, **A**) and sequential plaque area (**B**). **C.** Hematoxylin counterstain was used to identify necrotic cores. The percentage of total plaque area identified as necrotic is shown. **D.** Expression of the macrophage marker CD68 (red) and the smooth muscle cells marker αSMA (green) was determined by fluorescent immunohistochemistry. CD68 expression levels relative to plaque area and proportion of αSMA in fibrotic cap are shown. Representative image illustrating the staining is shown. **E.** Fibrotic cap (FC) thickness was defined as the shortest distance between the top of a necrotic core (*) and the lumen and the smallest measured FC for each animal is shown. Animals with plaques not displaying a visible necrotic core were excluded from the fibrotic cap thickness analysis. Circles denote individual animals; horizontal bars indicate the median value. *, *p*<0.05; **, *p*<0.01, indicated comparisons, Kruskall-Wallis (3 groups) or Mann-Whitney (2 groups) test. Tables indicate the effect of sex, nephropathy status and their interaction (two-way ANOVA), when the interaction was found significant. When significant, corresponding results are shown for males and females separately.

Consistent with the higher atherosclerotic burden seen in CKD^2–4^, AAN mice had more plaque occlusion at the aortic sinus than controls (Fig.1A, left). Notably, we observed a significant interaction between sex and nephropathy status on pooled plaque occlusion (*p*= 0.032), whereas the main effects of sex (*p*= 0.1443) and AAN status (*p=* 0.0653) alone were not significant (Fig.1A, right). This indicates that the impact of AAN on plaque size is restricted to females in this model. Subsequent post-hoc multiple comparisons showed that AAN significantly increased occlusion exclusively in female animals (Mean Difference = 11.34, 95% CI: 5.07-23.24). with no detectable effect observed in male animals. Sequential plaque area from the open valves was also increased in female AAN compared mice compared to controls (Fig.1B).

Plaque lipid content was not different between groups (not shown). In line with their increased size, plaques were more advanced in AAN females, as judged by a larger necrotic core area (Fig.1C).

In addition, AAN plaques in both sexes (no significant effect of sex observed) also displayed higher expression of CD68, indicating increased or more activated macrophage content (Fig.1D). Vascular smooth muscle cells (SMC, αSMA) distribution was found modestly altered in AAN with a trend to less SMC having migrated to the top of the plaque than in Control (Fig.1D). A thinner fibrous cap was confirmed by measurement of the smallest fibrous cap thickness AAN animals (Fig.1E) and indicates a more vulnerable plaque phenotype, likely more prone to rupture^23^. Thus, these alterations indicate a detrimental effect of nephropathy on plaque phenotype in both sexes.

### AAN leads to chronic inflammation and elevated circulatory DAMP levels. Several DAMPs are associated with future risk of CV events in CKD patients

Atherogenic lipid profiles were unaltered by chronic nephropathy: levels of total cholesterol, VLDL/LDL, and oxidised LDL (OxLDL) were elevated upon HFD but similar in control and AAN groups and levels of atheroprotective HDL were not different across groups (Fig.2A).

**Figure 2.**
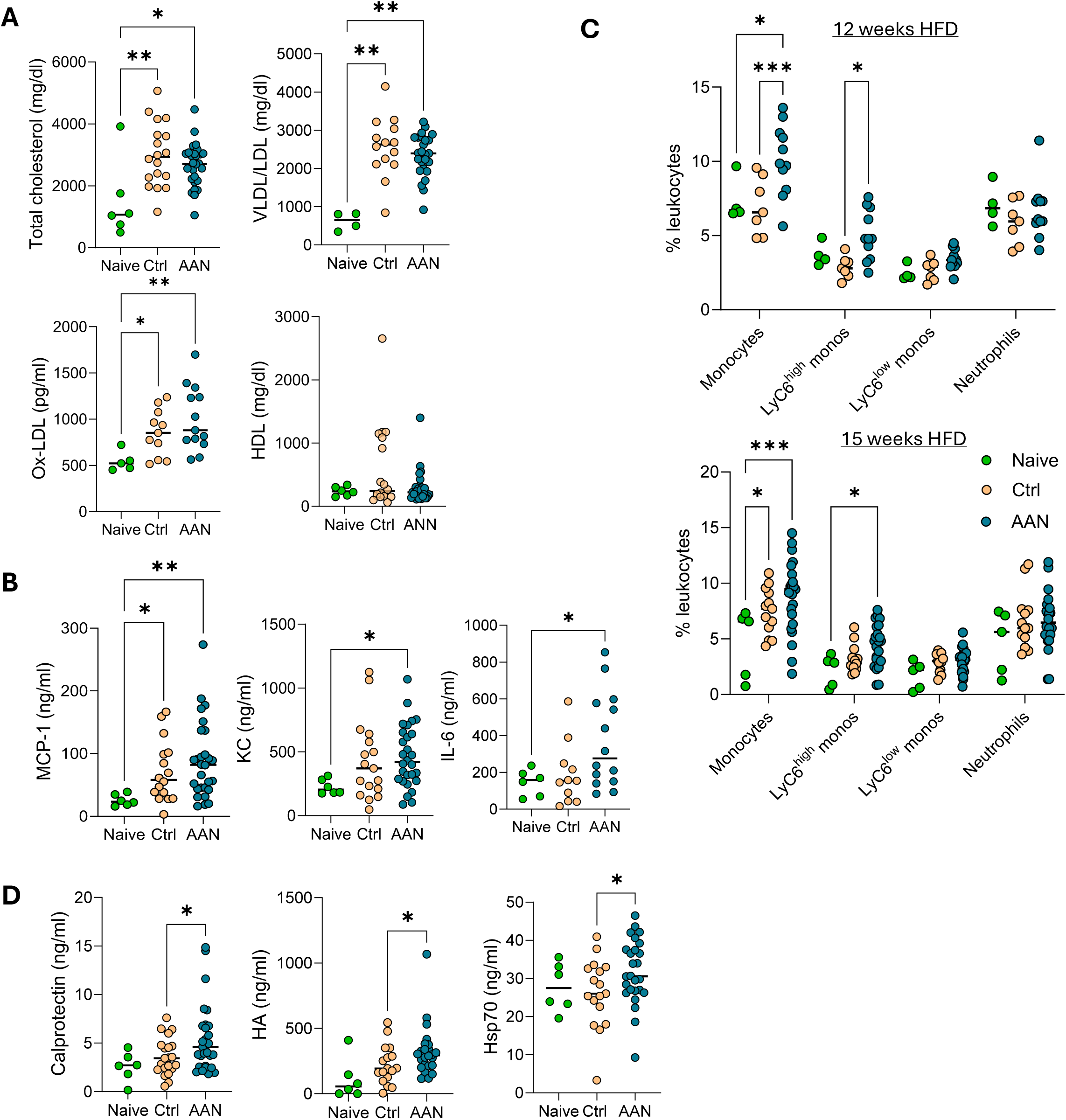
Limited changes to lipid profiles but increased chronic inflammation and circulatory. DAMP levels in *Ldlr ^-/-^* mice with AA-induced nephropathy *Ldlr ^-/-^* male and female mice were injected intraperitoneally 6 times at 2-day intervals with AA (3 mg/kg) or PBS (Control), starting on Day 0. On Day 12, normal chow diet was replaced with a high-fat diet (HFD). Blood was obtained by cardiac puncture after 15 weeks of HFD, unless stated otherwise. Age-matched *Ldlr ^-/-^* mice having not received injections or HFD served as a normal reference group (Naïve). Plasma lipid levels were determined by enzymatic reaction (**A**), cytokine and DAMP plasma levels by ELISA (**B, D**) and blood innate leukocyte proportions by flow cytometry (**C**). Circles denote individual animals; horizontal bars indicate the median value. *, *p*<0.05; **, *p*<0.01; ***, *p*<0.005, indicated comparisons, Kruskall-Wallis (3 groups) or Mann-Whitney (2 groups) test. No interaction between sex and nephropathy status were observed.

Conversely, the combination of AAN and HFD led to elevated plasma levels of pro-inflammatory cytokines (Fig.2B) compared to naïve mice. Monocyte proportions were also significantly elevated in AAN compared to control animals after 12 weeks of HFD (Fig.2C), an effect mostly driven by the pro-inflammatory Ly6C^high^ subset^24^, preferentially recruited to the atherosclerotic plaque^25^ and specifically associated with increased risk of CV events in CKD^15^. Total and Ly6C^high^ monocyte proportions remained elevated in AAN mice after 15 weeks of HFD, although the increase compared to control animals was no longer significant due to an increase in monocyte proportions in response to HFD at this time point (Fig.2C).

Together, these data suggest that exacerbation and alteration of atherosclerosis in AAN mice are driven by systemic inflammation, rather than by a more atherogenic lipid profile.

Circulatory levels of several DAMPs were found elevated in CKD patients, notably Calprotectin, HA, Hsp70^11^ and Hsp90^26^. Here, levels of these DAMPs were also raised in plasma from AAN animals (Fig.2D).

Supporting a role of elevated DAMPs in promoting CV risk in CKD, both subunits of Calprotectin (S100A8/S100A9) were reported among the top elevated proteins in CKD3/4 patients who went on to suffer from a CV event (n=32), compared to matched patients who did not (n=34)^17^. However, careful repurposing of the data showed that several other TLR DAMPS are also significantly elevated in plasma from CKD patients who later experienced a CV event, including Histones and Heat-shock proteins (Table 1).

**Table 1.**
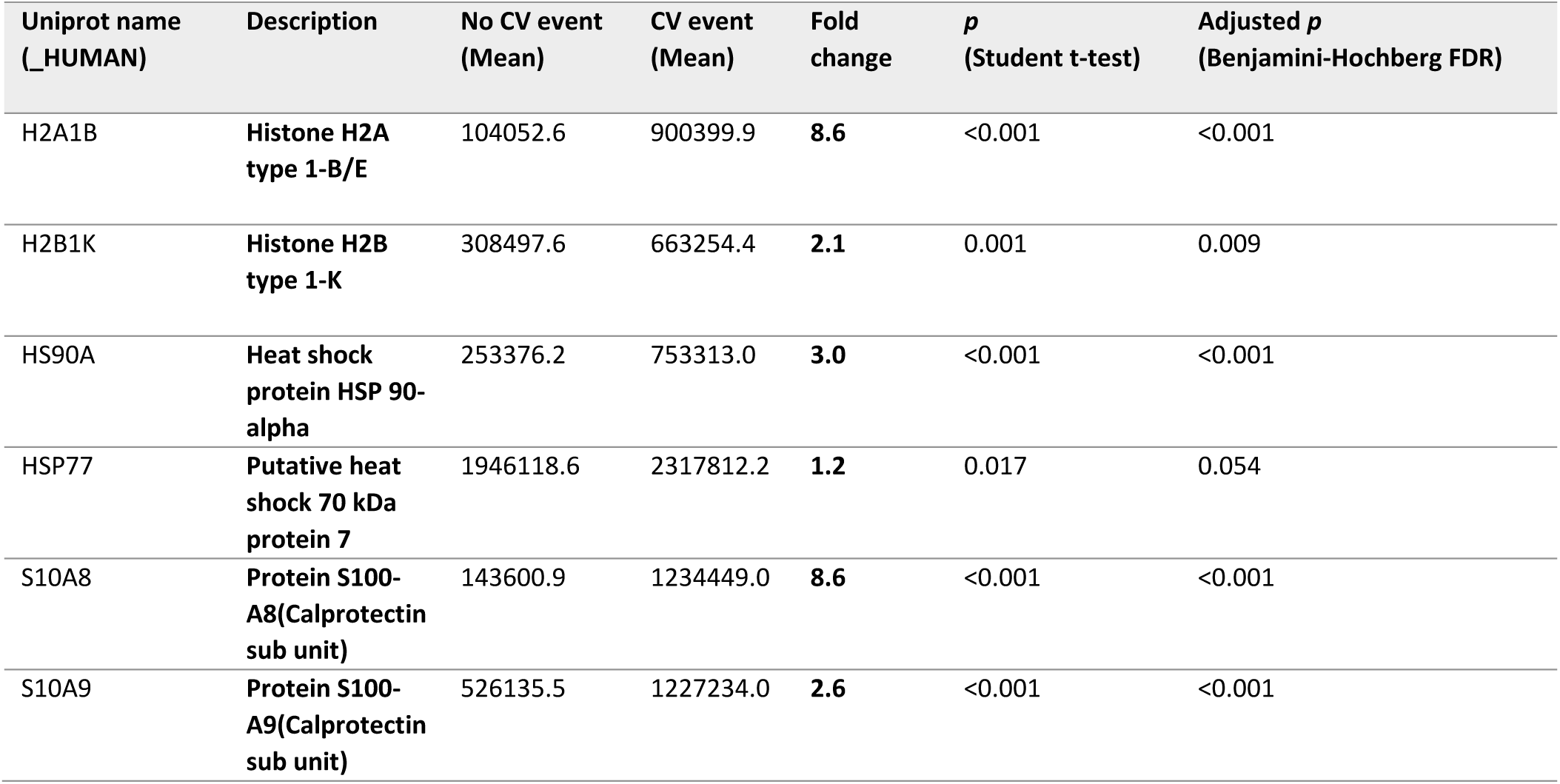
Known TLR DAMPs found at higher levels in CKD patients who subsequently underwent a CV event (=32), compared to those who did not (n=34)

### Single-cell transcriptomic analysis highlights increased TLR-related inflammatory pathways in aortas from AAN mice

To further investigate the pathways that promote atherosclerosis in CKD, a comparative single-cell transcriptomic analysis of aortas in control and AAN mice was performed. We applied unbiased Louvain clustering on 83,456 cells, identifying 16 clusters (Fig.3A, Sup. Fig.2A), which were assigned a cell type based on established lineage markers (Sup. Fig.2B). We observed 9 nonimmune clusters, including 3x vascular SMCs (Clusters 0, 7, 11), differentially expressing *Notch3* (higher in Cluster 7, Notch3 SMC), and *Col3a1* and *Mgp* (SMC-1 > SMC-2); 3x fibroblasts (Clusters 3, 13, 14, Fibroblasts-1 to 3), differentially expressing *Lum*; 2x endothelial cells (EC, Clusters 1 and 10), differentially expressing *Cd34* (lower in Cluster 10, Activated EC) and 1x Mesenchymal cells. Seven immune cell clusters were found, including 3x T-cells (Clusters 4, 6, 8), 3x B-cells (Clusters 2, 12, 15) and 1x Macrophages (Cluster 5).

**Figure 3.**
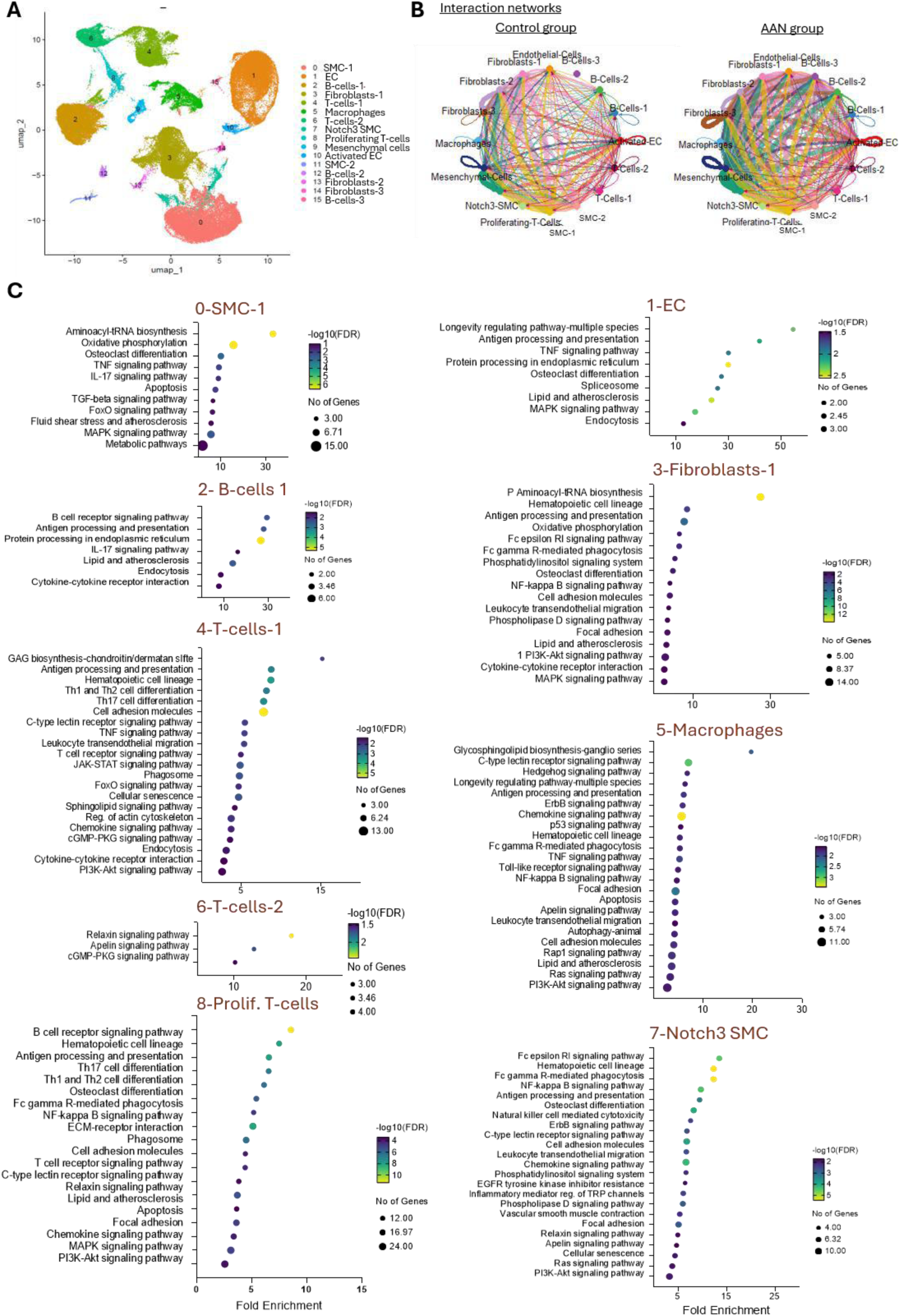
Single cell transcriptomic analysis reveals an enrichment in inflammatory and atherosclerotic disease pathway in AAN vs Control aortas. Single cells were isolated from murine aortas (n=6 animals/experimental group, 3 females, 3 males, biological replicates for A-C) before undergoing full single-cell transcriptomic analysis (scRNAseq) **A.** UMAP projection of cells (n=83,456 cells) highlights the 16 identified cell clusters **B.** CellChat circle plots show inferred intercellular communication in-between cell clusters. Node size is proportional to cell population size, arrow width reflects the strength of predicted ligand-receptor interactions, and arrow colour indicates signal sender cell**. C.** Graphs display the upregulated KEGG pathways in AAN vs Control (Ctrl) mice for each of the Louvain clusters. False Discovery Rates (FDR) were calculated by adjusting raw *p*-values from a hypergeometric distribution test using the Benjamini-Hochberg method for each pathway and pathways were ranked by lowest enrichment FDR. Pathways containing irrelevant terms were removed. Clusters not shown in (C) did not exhibit AAN-upregulated KEGG pathways.

T-cells clusters differentially expressed *mKi67* (higher in Cluster 8, Proliferating T-cells) and *Rorc* (higher in Cluster 6, T-cells-2, suggesting an enrichment in Th17 T-cells, Sup. Fig.2B). These clusters and populations are in line with those described by others in human and mouse atherosclerosis^27^, although these studies did not incorporate nephropathy.

A CellChat analysis revealed a complex and dense intercellular communication network (Fig.3B), as previously shown^27^. The profile was consistent with stromal and vascular cells as major signalling hubs and immune populations mostly as recipients of signals. Interestingly, AAN doubled the number of cellular interactions inferred (Sup. Fig.3C), strengthening non-immune to immune cell communications, as well as non-immune to non-immune (Fig.3B).

KEGG pathway analysis on a mixed-sex population identified pathways significantly upregulated by AAN in the 9 most abundant clusters (Clusters 0 to 8, Fig.3C). The *Lipid and atherosclerosis* pathway was enriched in several AAN clusters. Canonical pathways associated with TLR/PRR activation were also among the top upregulated pathways in many clusters (Fig.3C), supporting our underlying hypothesis that the DAMP:TLR pathway plays a critical role in promoting atherosclerosis in CKD. These included *NF-kappa B signaling pathway* (Fibroblasts 1, Notch3 SMC, Macrophages and Proliferating T-cells), *MAPK signaling pathway* (SMC-1, EC, Fibroblasts-1 and Proliferating T-cells) and *TNF signaling pathway* (SMC-1, EC, T-cells-1, Macrophages). Macrophages in AAN plaque also overexpressed the *TLR signaling pathway*, consistent with their high TLR expression. *C-type lectin receptor signaling,* another PRR system responding to DAMPs, was also upregulated in Macrophages, as well as in Notch3 SMCs and Proliferating T-cells.

Other AAN-upregulated pathways consistent with (although not specific of) increased DAMP:TLR signaling included pathways associated with i) immune cell adhesion and recruitment: *Cell adhesion molecules* (Fibroblasts-1, Notch3 SMC, Macrophages, Proliferating T-cells), *Leukocytes transendothelial migration* (Fibroblasts-1, Macrophages), *Focal adhesion* (Fibroblasts 1, Notch3 SMC, Macrophages, Proliferating T-cells), *Rap1 signaling pathway* (Macrophages), *ECM-receptor interaction* (Proliferating T-cells) ; ii) response to cytokines: *Cytokine-cytokine receptor interaction* (B-cells-1, Fibroblasts-1, T-cells-1), *Chemokine signaling pathway* (T-cells-1, Notch3 SMC, Proliferating T-cells); iii) cell death: *Apoptosis* (SMC-1, Macrophages, Proliferating T-cells), *Cellular senescence* (T-cells-1, Notch3 SMC), *p53 signaling* pathway (Macrophages) ; and iv) *Antigen processing and presentation* (EC-1, B-cells-1, Fibroblasts-1, Macrophages, Proliferating T-cells).

AAN also promoted a range of cell-specific atherosclerotic responses, such as *TGFβ signaling* in SMC-1, *Endocytosis* in EC, *B-cell receptor signaling* in B-cells 1, *T-cell differentiation* and *T-cell receptor signaling* in T-cells 1 and Proliferating T-cells, *Fc gamma receptor mediated phagocytosis* in Macrophages. Both SMC clusters also upregulated the *Osteoclast differentiation pathway*, suggesting a possible phenotype switch that will promote matrix degradation ^28^, in line with the thinner fibrous caps observed in AAN mice, and expected to promote plaque instability.

Therefore, our transcriptomic findings support a role of inflammatory pathways in driving exacerbated atherosclerosis in AAN mice and are consistent with enhanced TLR activity.

Given the sex-differential effect of AAN on plaque size and necrotic core area, transcriptomic changes were also analysed in male and female aortas separately. Clusters similar to those described for both sexes together were identified in both sexes (Sup. Fig.3A-C), although a lower number of clusters were identified in females, probably because of a lower cell number (Males, 59977; Females, 23479). Cluster proportions were similar across sexes (Sup. Fig.3D), except for T-cells clusters (-2 and proliferating), which were noticeably larger in males (proliferating T cells did not cluster separately in females). For some clusters (e.g. Macrophages), less DEGs and less/no enriched pathways were identified within either sex, likely because of lower cell numbers per cluster and, thus, reduced power. However, many of the pathways identified above were also found enriched in AAN males and/or female clusters (Sup. Tables 1 to 12), centred around general disease pathways (purple), immune cell adhesion and recruitment (yellow), inflammation (red), cell death (green) and cell-specific pro-atherosclerotic responses (blue).

### Calprotectin inhibition only modestly ameliorates atherosclerosis in AAN mice

Although Calprotectin was identified as the DAMP with the strongest association with future CV events in CKD patients^17^ and was also elevated in AAN mice, we found that Calprotectin blockade, initiated at Day 29 using its well-described pharmacological inhibitor Paquinimod at a preestablished dose^17^, did not lead to a lower atherosclerosis burden (Fig.4A,B). Its beneficial effect on plaque phenotype was restricted to driving a thicker fibrous cap (Fig.4F), without improving necrotic core or macrophage/SMC plaque composition (Fig.4C-E). This is despite our previous findings that Calprotectin can promote several key atherosclerotic responses *in vitro*^11^. We performed further analysis of our data to rank DAMPs according to their ability to modulate key atherosclerotic functions, highlighting that Calprotectin was a potent disruptor of trans-endothelial resistance and promoter of foam cell formation. However, it was a poor inducer of MCP-1 production and promoter of migration in monocytes (Sup. Fig.4). Of note, most pro-atherosclerotic responses tested were promoted by at least 2 of the 3 DAMPs tested, indicating redundancy of function. Thus, while we cannot exclude that inhibition of a DAMP other than Calprotectin may be more efficient at reducing atherosclerosis in CKD, these data support the concept that multiple DAMPs may contribute to exacerbated atherosclerosis in CKD and that multi-DAMP targeting may be required to achieve robust therapeutic effects.

**Figure 4.**
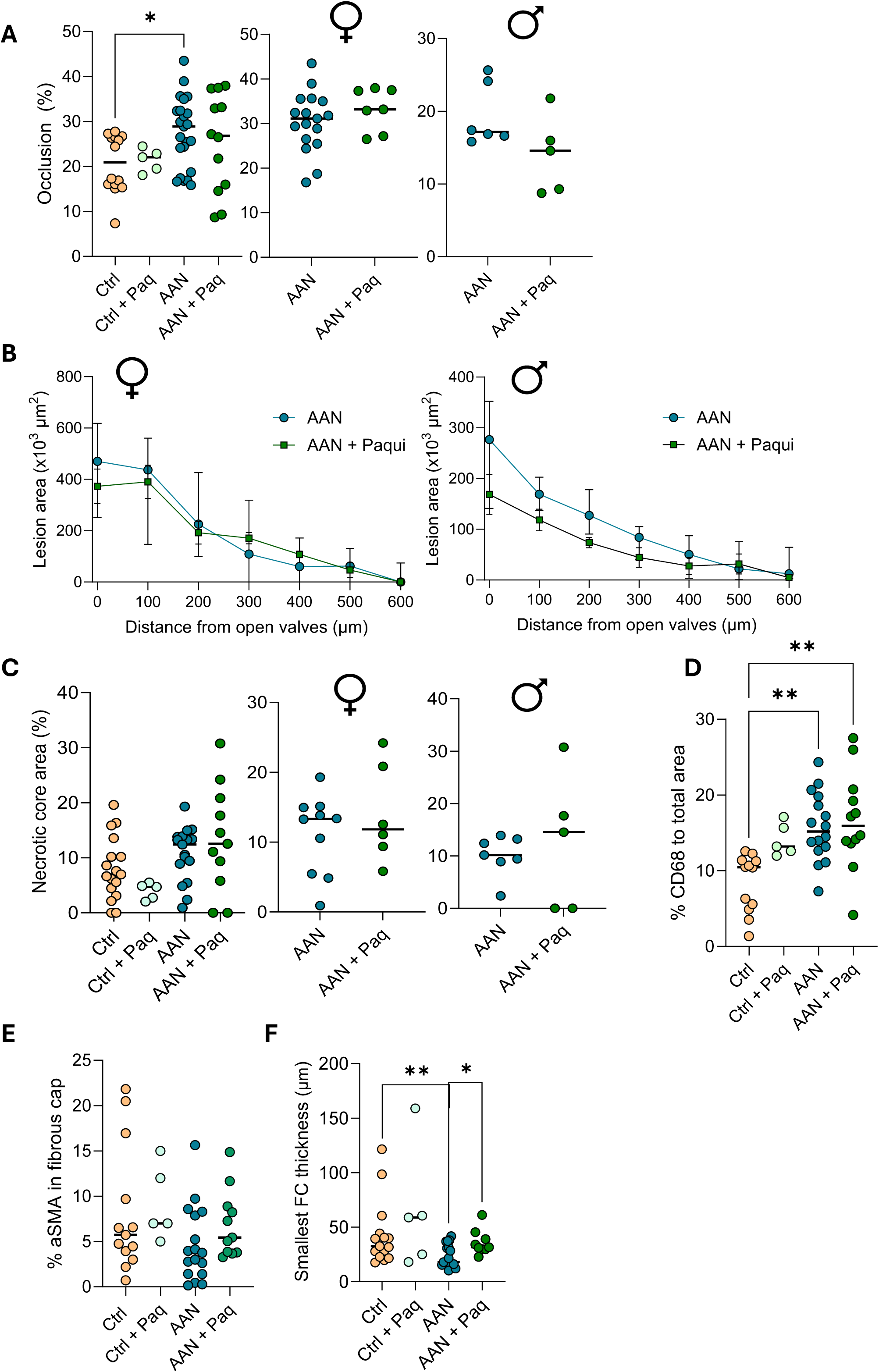
Calprotectin blockade only modestly ameliorate atherosclerotic disease in AAN mice. *Ldlr ^-/-^* male and female mice were injected intraperitoneally 6 times at 2-day intervals with AA (3 mg/kg, AAN) or PBS (Ctrl), starting on Day 0. On Day 12, normal chow diet was replaced with a high-fat diet (HFD). At Day 29, after kidney disease is established in this model [11], daily oral administration of Calprotectin inhibitor Paquinimod (Paq) in drinking water (5mg/kg/day^17^) was initiated. Experiments were terminated after 15 weeks of HFD. Frozen sections of the aortic root were obtained and sections displaying the 3 valves (A, C-F) or sequential sections (B) were analysed for atherosclerosis. **A-B.** Oil Red O staining was used to determine the percentage of occlusion (plaque to lumen area, **A**) and sequential plaque area (**B**). **C.** Hematoxylin counterstain was used to identify necrotic cores and the percentage of total plaque area identified as necrotic is shown. **D-E.** Expression of the macrophage marker CD68 (**D**) and the smooth muscle cells marker αSMA (**E**) was determined by fluorescent immunohistochemistry. **F.** Fibrotic cap (FC) thickness was defined as the shortest distance between the top of a necrotic core and the lumen and the smallest measured FC for each animal is shown. Circles denote individual animals; horizontal bars indicate the median value. *, *p*<0.05; **, *p*<0.01, indicated comparisons, Kruskall-Wallis (3 groups) or Mann-Whitney (2 groups) test. Results in (A-C) are shown for males and females separately, as the effect of nephropathy status was found to be sex-dependent. Sex did not influence the lack of effect of Paquinimod treatment.

### Soluble TLR2 reduces atherosclerotic plaque development and improves plaque stability in AAN mice

sTLR2 is a naturally occurring inhibitor of TLR activity that can reduce activation of multiple TLRs by binding to and reducing the enhancing activity of the common TLR coreceptor CD14^16,29^. Weekly administration of sTLR2 was initiated once nephropathy is established (Day 29^11,14^, Sup. Fig.5A) and administration of sTLR2 did not lead to a reduction in kidney pathology (Sup. Fig.5B,C) or alter lipid profiles (Sup. Fig.5D). However, sTLR2 significantly reduced AAN-associated chronic inflammation, as judged by lower plasma cytokine levels (Fig.5A) and monocyte proportions (Fig.5B). Atherosclerotic disease in AAN animals was also significantly ameliorated by sTLR2 treatment (Fig.5C-G). Specifically, AAN mice treated with sTLR2 displayed a substantially lower atherosclerotic burden, comparable to that of controls (Fig.5C, D). sTLR2 also promoted a less vulnerable and inflammatory plaque phenotype, as judged by smaller necrotic core size (Fig.5E), a thicker fibrous cap (Fig.5F), together with more SMC at the cap and reduced plaque CD68 expression (Fig.5G). Of note, sTLR2 did not lower inflammation or atherosclerosis in control mice, indicating its specific effect on AAN-mediated inflammation.

**Figure 5.**
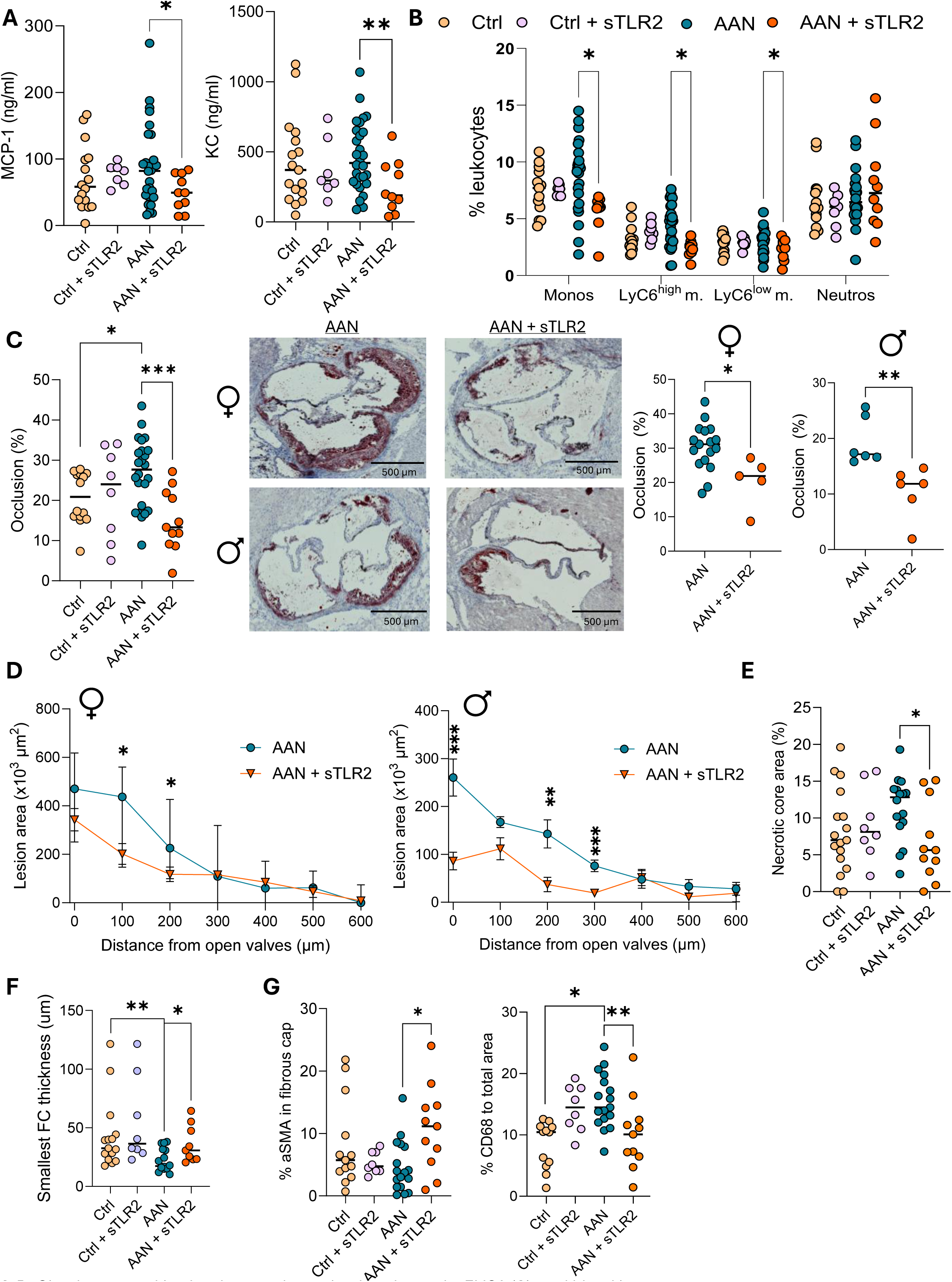
Treatment with sTLR2 prevents the exacerbating impact of AAN on atherosclerosis. *Ldlr ^-/-^* male and female mice were injected intraperitoneally 6 times at 2-day intervals with AA (3 mg/kg, AAN) or PBS (Control), starting on Day 0. On Day 12, normal chow diet was replaced with a high-fat diet (HFD) for 15 weeks. On Day 29, after kidney disease is established in this model [11], weekly i.p. administration of sTLR2 (12.5 µg/kg) was initiated in the treatment and control groups, until the end of the experiment. **A-B.** Circulatory cytokine levels were determined in plasma by ELISA (**A**), and blood innate leukocyte proportions by flow cytometry (**B**). **C-G.** Frozen sections of the aortic root were obtained and sections displaying the 3 valves (C, E-G) or sequential sections (D) were analysed for atherosclerosis. **C-D**. Oil Red O staining was used to determine the percentage of occlusion (plaque to lumen area, representative images shown, **C**) and sequential plaque area (**D**). **E.** Hematoxylin counterstain was used to identify necrotic cores. The percentage of total plaque area identified as necrotic is shown. **F**. The smallest measured fibrotic cap (FC) thickness for each animal is shown. **G.** Macrophages (CD68, red) and smooth muscle cells (αSMA, green) were detected by fluorescent immunohistochemistry. CD68 signal relative to plaque area, and αSMA distribution to the fibrous cap are shown. Circles denote individual animals; horizontal bars indicate the median value. *, *p*<0.05; **, *p*<0.01; ***, *p*<0.005, indicated comparisons, Kruskall-Wallis (3 groups) or Mann Whitney (2 groups) test. Results in (C-D) are shown for males and females separately, as the effect of nephropathy status was found to be sex-dependent. Sex did not influence the effect of sTLR2 treatment.

The effect of the sTLR2 in restricting and stabilising atherosclerotic plaques in AAN animals confirms the critical role of TLR agonists in driving these effects. Because CKD may increase gut permeability^30^, leading to endotoxin entering the bloodstream^31^ and activating systemic TLRs, we verified that endotoxin levels in Control and AAN groups were similar (Ctrl: 0.00356 ng/ml +/-0.0002 vs AAN: 0.00348 ng/ml +/-0.0001, *p*=0.756). We therefore conclude that AAN animals do not display increased gut permeability and thus that increased microbial stimulation of TLRs cannot account for the worsening effect of AAN on atherosclerosis. Taken together, our data indicates that the increased plaque burden and more vulnerable phenotype observed in AAN animals is the result of TLR engagement by DAMPs.

Consistent with sTLR2’s anti-inflammatory mode of action, scRNAseq analysis revealed that its administration inhibited many of the inflammation-driven pathways that were found enriched in AAN aortas (Fig.6A). Notch3 SMCs and Macrophages were the clusters most affected by this effect, as sTLR2 downregulated 30% and 32% (respectively) of AAN-upregulated pathways in these cells (Fig.6B). Of particular relevance, sTLR2 downregulated Toll*-like receptor signaling* in Macrophages, *NF-kB signaling* in Notch3 SMCs and both clusters downregulated *Antigen processing and presentation*, in line with the TLR control of adaptive immunity. *Osteaclast differentiation* was also downregulated in Notch3 SMCs and SMC-1, consistent with the observed increased fibrotic cap thickness.

**Figure 6.**
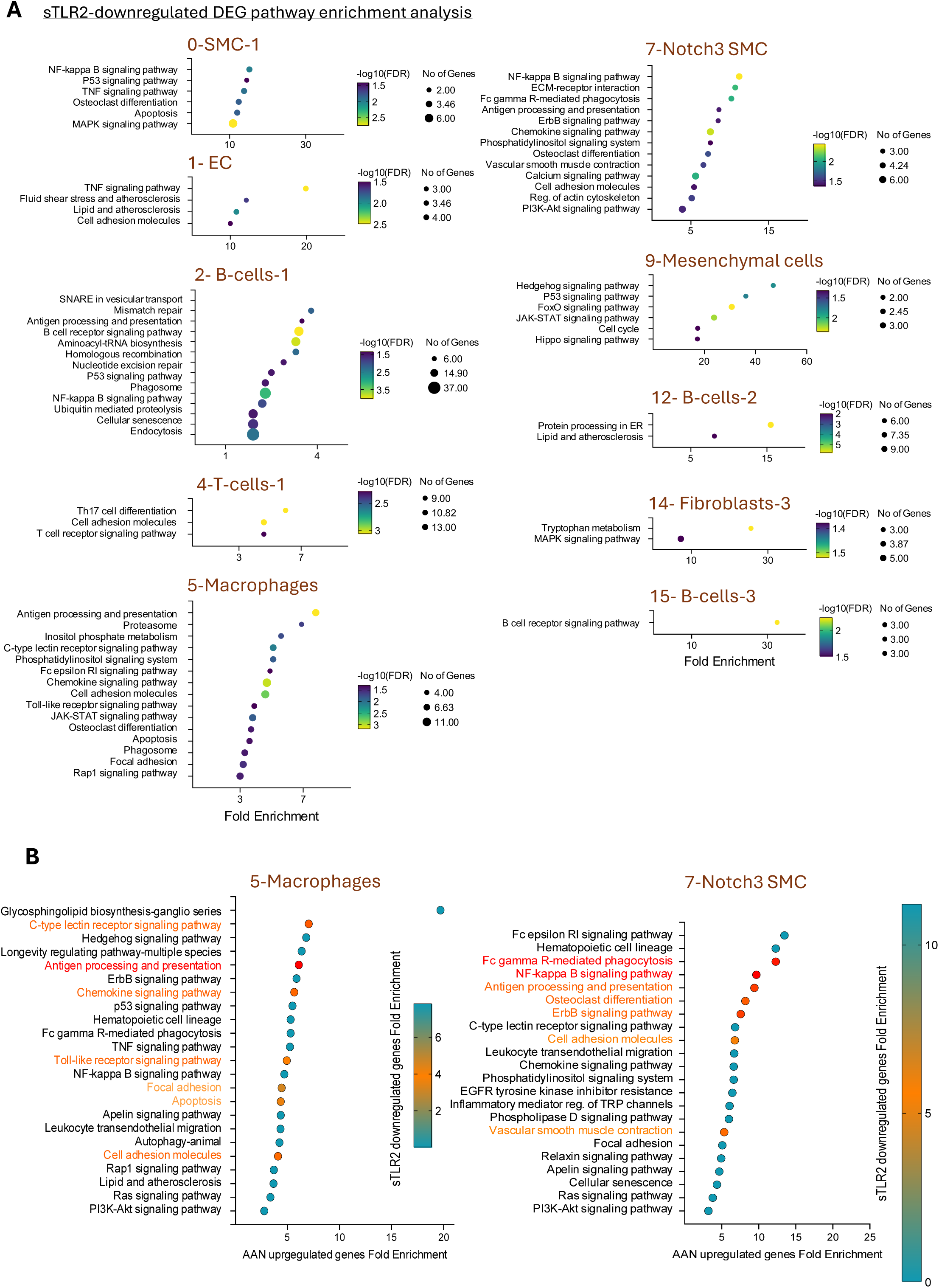
Single cell transcriptomic analysis highlights a wide-ranging inhibition of inflammatory and atherosclerotic disease pathways in sTLR2 treated AAN mice. Single cells were isolated from murine aortas (n=6 animals/experimental group, 3 females, 3 males) before undergoing full single-cell transcriptomic analysis (scRNAseq). A total of 83,456 cells were analysed. **A.** Graphs display the KEGG pathways enriched in DEGs downregulated by sTLR2 treatment (AAN + sTLR2 vs AAN groups) for each of the Louvain clusters. False Discovery Rates (FDR) were calculated by adjusting raw *p*-values from a hypergeometric distribution test using the Benjamini-Hochberg method for each pathway and pathways were ranked by lowest enrichment FDR. Pathways containing irrelevant terms were removed. Clusters not shown did not exhibit sTLR2-downregulated KEGG pathways. **B.** Graphs display the AAN-upregulated KEGG for the indicated cell clusters (AAN vs Ctrl), and colour-coding indicates the strength of sTLR2 inhibition of these pathways, measured by enrichment in sTLR2 downregulated genes (AAN + sTLR2 vs AAN).

## DISCUSSION

CKD leads to markedly increased cardiovascular risk^2,3,32^ and CV treatments effective in the non-CKD population offer less protection in CKD^32^. The unique state of chronic inflammation in CKD is thought to drive the suboptimal responses to lipid-lowering or blood pressure-reducing therapies seen in these patients. Trials using antibodies to IL-1β demonstrated that reducing chronic inflammation reduces CV risk in CKD patients but carried an increased risk of severe infections leading to death^33,34^. Similarly, low-dose Colchicine, a potent anti-inflammatory that notably blocks inflammasome activation, was found efficient at reducing severe CV events in patients with prior myocardial infarction, but doubled the risk of pneumonia^35^. Thus, the development of safe strategies to reduce CV risk in CKD remains a priority.

By interfering with the activity of CD14, a TLR co-receptor which is required to varying extent by TLR ligands, sTLR2’s provides partial - but typically not complete - inhibition of TLR activation. Consequently, sTLR2 administration does not compromise bacterial clearance, in normal^16^ or nephropathic^11^ mice. This distinctive mode of action offers a unique opportunity to dampen the activity of multiple DAMPs while preserving sufficient pathogen recognition. This is especially relevant as we highlight that several DAMPs are elevated in CKD patients who subsequently experience CV events and demonstrate that several DAMPs can drive atherosclerotic responses *in vitro*. This suggests the contribution of multiple DAMPs to the pathology, in line with our observation that the Calprotectin inhibitor Paquinimod only modestly ameliorates atherosclerotic disease in nephropathic mice.

Interestingly, and despite TLR involvement in atherosclerosis^8,9^, sTLR2 did not reduce plaque burden in non-AAN animals, indicating that its effect is mediated by reducing kidney disease-associated inflammation specifically. This is despite OxLDL, the major initiator of atherosclerosis, being a well-described TLR4 and TLR2 agonist. STLR2’s lack of effect on atherosclerosis in the absence of AAN may have several explanations. Firstly, the regimen of weekly sTLR2 administration chosen here may be insufficient to inhibit the activity of the high levels of atherogenic lipids induced by HFD. Secondly, the requirement for CD14, sTLR2’s main target, to facilitate OxLDL recognition by TLRs is not described. It is possible that OxLDL activation of TLRs may happen without/with minimal requirement for CD14.

Our study does not address the origin of elevated systemic DAMPs in CKD. It is likely that both ongoing kidney inflammation and reduced renal clearance will promote the systemic accumulation of inflammatory mediators and toxins, thus leading to DAMP release from blood cells or the endothelium. In addition, there is ample evidence of elevated renal DAMPs in CKD^36^, which may reach the bloodstream. Here, and although the last insult to the kidneys occurred 4 months earlier, we found that levels of MCP-1 and HA remained elevated in AAN kidneys at the end of the experiment (MCP-1: Ctrl, 81.8 pg/mg protein+/-7.8 *vs* AAN, 98.5+/- 4.8, *p=*0.0077; HA: Ctrl, 52.14 ng/mg protein+/-5.9 *vs* AAN, 70.10+/-4.2, *p=*0.0261). Supporting a role for the inflamed kidney mass in promoting exacerbated atherosclerosis in CKD, 5/6 nephrectomy did not lead to increased atherosclerotic burden in ApoE^-/-^ mice^37^. Therefore, models of kidney removal were not used here.

A drawback of most common CKD models for the study of atherosclerosis is reduced food intake^38^. We observed that AAN males, but not females, had lower body weights than their control counterparts at the end of the experiment (Ctrl: 33.8 g+/-1.351 vs AAN: 29.8 g +/- 1.167, *p*=0.038). Thus, we conclude that the lack of increase in atherosclerotic plaque size in AAN vs Control males may be due to reduced HFD intake by AAN male mice, rather than a genuine lack of effect of AAN on atherosclerotic plaque size in males. Nonetheless, AAN male mice displayed a significantly more inflammatory and vulnerable plaque phenotype than controls, together with a profound enrichment in inflammatory pathways. Furthermore, treatment with sTLR2 led to a reduction in atherosclerosis plaque size in AAN male mice, confirming the ability of AAN-induced DAMPs to drive plaque atherosclerotic plaque progression in males.

Of note, this is the first report of single-cell transcriptomic analysis of atherosclerotic arteries in chronic nephropathy (mouse or human). This analysis revealed that chronic nephropathy dramatically increased cell:cell communication between clusters and promoted inflammatory pathways in most cell types identified. This supports a critical role of inflammation in promoting atherosclerosis in CKD and is consistent with the involvement of broadly expressed innate immune pathways in amplifying vascular inflammation and accelerating atherosclerotic disease.

Together with increased plaque burden, medial vascular calcification is a significant driver of CV risk in CKD^2,32^. Calprotectin was found to mediate plaque calcification in CKD^17^, although it is unclear whether it drives the more harmful medial calcification. Whether sTLR2 can mitigate calcification remains unclear, as our model did not exhibit this pathology, although sTLR2 did modulate vascular SMC phenotype.

Thus, this study establishes the basis for translational investigations into the potential of sTLR2 to improve cardiovascular outcomes in the challenging CKD population. Our experimental approach targets inflammation occurring after CKD is established, highlighting its potential for translation to therapy and allowing personalised targeting of high-CV risk patients.

## SUPPLEMENTARY MATERIAL CONTENT

Supplementary Figures 1 to 5

Supplementary Tables 1 to 12

Supplementary Methods

